# Long-range Hill-Robertson effect in adapting populations with recombination and standing variation

**DOI:** 10.1101/2022.11.07.515399

**Authors:** Igor M. Rouzine

## Abstract

In sexual populations, closely-situated genes have linked evolutionary fates, while genes spaced far in genome are commonly thought to evolve independently due to recombination. In the case where evolution depends essentially on supply of new mutations, this assumption has been confirmed by mathematical modeling. Here I examine it in the case of pre-existing genetic variation, where mutation is not important. A haploid population with *N* genomes, *L* loci, a fixed selection coefficient, and a small initial frequency of beneficial alleles *f*_0_ is simulated by a Monte-Carlo algorithm. The results demonstrate the existence of extremely strong linkage effects, including clonal interference and genetic background effects, that depend neither on the distance between loci nor on the average number of recombination crossovers. When the number of loci, *L*, is larger than 4log^2^(*Nf*_0_), beneficial alleles become extinct at most loci. The substitution rate varies broadly between loci, with the fastest rate exceeding the one-locus model prediction. All observables and the transition to the independent-locus limit are controlled by single composite parameter log^2^(*Nf*_0_)/*L*. The potential link between these findings and the emergence of new Variants of Concern of SARS CoV-2 is discussed.

## Introduction

A typical species is heterozygous at millions of genomic sites, loci. The average difference between an individual’s genome and the consensus genome is estimated at 20 million base pairs, or 0.6% of the total of 3.2 billion base pairs (1). The invention of the new methods of full-genome DNA sequencing caused the emergence of the field of genomics and proteomics dedicated to the quantitative aspects of genetic diversity and gene expression at a large number of loci (2–7). To describe and visualize the genetic complexity, various computational methods have been developed including phylogenetics, the principle-components analysis, the cluster analysis. Among them, mathematical modeling of evolution stands out as a tool of a high predictive power. Modeling allows to connect, in the most direct and reproducible fashion, the assumptions about the dominant factors of evolution to the predictions for the observable parameters of genetic diversity and evolutionary dynamics.

The assumptions and simplifications of models vary broadly depending on the systems studied and the questions asked. Two distinct groups of models and methods have been applied to animal populations and microbial populations. The classical one-locus and two-locus models that neglect interaction with the other loci in genome (8–10) dominate the way in which many evolutionary biologists think about the evolution of higher organisms. In contrast, monocellular eukaryotes, viruses, and bacteria that are characterized by an extremely high genetic diversity and ultrarapid evolution, are often described by asexual or partly sexual population models that include explicitly large numbers of interacting loci. Analysis of the evolutionary dynamics of multi-locus models is more complex than one-locus and two-locus models and relies either on Monte-Carlo simulation (11–17) or the advanced mathematical methods of statistical physics (11, 18–39). The heavy mathematical artillery is required, because the evolution of many different loci is inter-dependent (40). There are two kinds of interference effects. One kind, not considered in this article, is epistasis arising from biological interaction of different loci, including protein-protein interactions or interactions gene regulation network (29, 31–33, 41–52). The second type, which is the focus of the present article, is the effects originating from the common ancestry of different loci, including Hill–Robertson effect, clonal interference, background selection and hitchhiking (8, 25, 53–55). Linkage effects also slow down adaptation (11, 21, 23, 26), increase accumulation of deleterious alleles (11, 21), and change the statistical shape of genealogical tree (24, 27, 56).

In sexually reproducing organisms and organisms with frequent recombination such as some viruses, linkage effects are partly compensated by recombination between parental genomes. A fundamental fact of genetics discovered by Morgan is that frequent recombination destroys allelic associations, so that alleles at far-spaced loci segregate independently. Conventional wisdom tells us that all the other linkage effects between far-situated loci must vanish as well. A model of long-term sexual evolution limited by rare mutation seemed to confirm this expectation (57). Assuming that genome consists from independently-evolving blocks and applying the phylogenetic theory of asexual evolution to each block, the authors constructed a scaling argument expressing the length of each block, the lead of the traveling wave, and the average coalescent time in terms of the average adaptation rate. The analytic predictions have been confirmed numerically for two particular models of population in the presence of natural selection and mutation.

In the present work, I investigate linkage effects in a different biological scenario, when natural selection and recombination act on pre-existing beneficial alleles, and new mutations can be neglected. This model is appropriate in the case when selection pressure changes its sign at a large number of loci. For example, a population migrates to a new environment, or a virus is subjected to the immune response or a replication inhibitor treatment. In this case, weakly deleterious alleles pre-existing in the mutation-selection balance can become beneficial.

## Results

### Model

Consider a sexually reproducing population comprised of *N* individual genomes (or *N*/2 diploid genomes without allelic dominance), where each genome has *L* loci. In the beginning, each locus is assumed to have a fraction *f*_0_ of beneficial alleles, with fitness benefit s. The value of *f*_0_ is assumed to be in interval 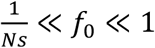. Next, I assume that a genome undergoes an average number *M* of random crossovers with another, randomly chosen genome, and one of the two parents is replaced with the recombinant. The evolution is simulated using a Wright-Fisher process, in which the progeny genomes replace the parental genome, and the average progeny number is proportional to the genome fitness. The evolutionary factors included in the model are directional natural selection, random genetic drift, linkage, and recombination. New mutation and epistasis are absent. The details of simulation are described in the *Methods* section.

### Extinction of beneficial alleles depends on a single composite parameter

If the number of loci *L* is sufficiently large, beneficial alleles at most loci become extinct. The fraction of remaining polymorphous loci, denoted 1 – *C_loss_*(*t*), decreases in time from 1 to at a low plateau (Fig. 1A, red line). This result differs from the prediction of the single-locus model, in which multiple lineages per site are expected to reach fixation at *Nf*_0_*s* ≫ 1. In that case, the fixation probability of an allele is *s*, and the extinction probability is 1 – *s* (40). The probability of the extinction of all *Nf*_0_ beneficial lineages is given by *C_loss_*(∞) = (1 – *s*)^*Nf*_0_^ ≈ *e*^*Nf*_0_*s*^, which is exponentially small.

**Fig. 1.**
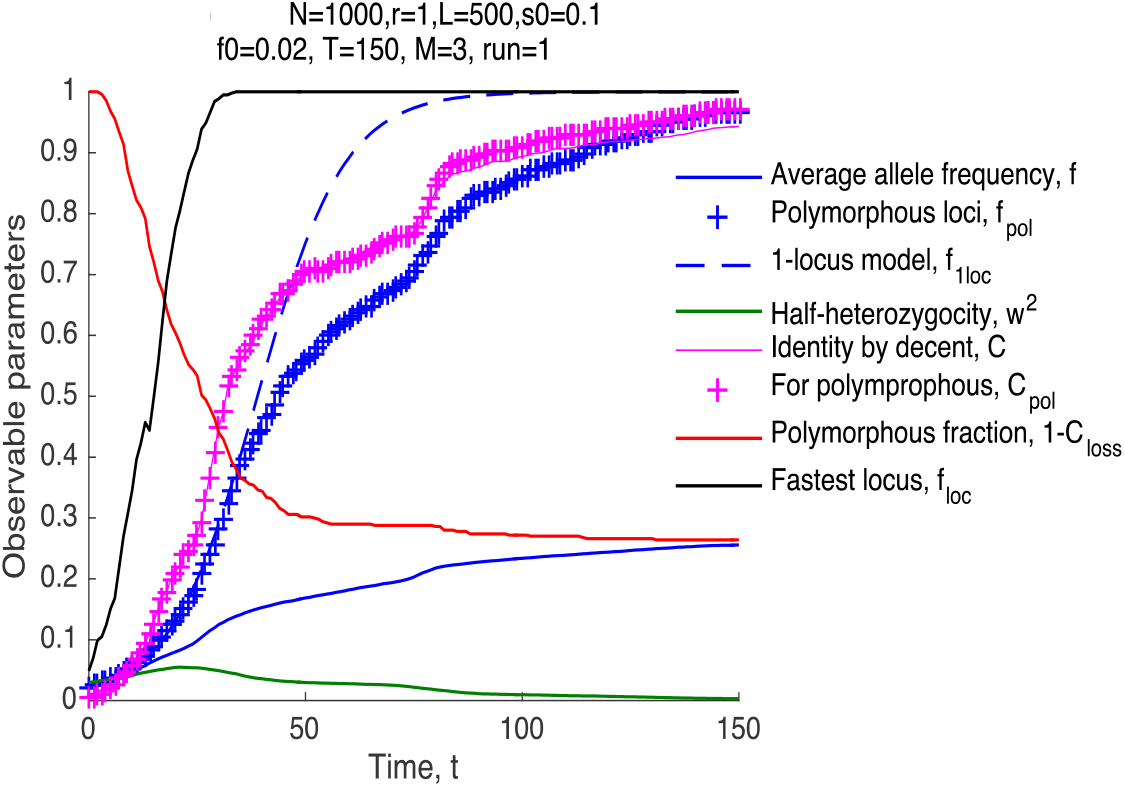
Dynamics of observables in the model with standing variation and the absence of mutation. Beneficial alleles become extinct at most loci. X-axis: Time in generations, *t*. Y-axis: Observable parameters calculated during simulation. The average frequency of beneficial alleles per locus per individual, *f*, the same value averaged over polymorphous loci only, *f_pol_*, the prediction for *f* of the deterministic one-locus model, *f*_1*loc*_, half-heterozygocity *w*^2^ = 〈*f*(1 – *f*)〉, the fraction of homologous pairs of loci with a common initial ancestor, *C*, the same value for polymorphous loci, *C_pol_*, the fraction of polymorphous loci, 1 – *C_loss_*, and the largest of allelic frequencies among loci, max (*f_loc_*). Parameter values are shown on the top. Parameters are defined in *Methods* and values are shown.

Varying model parameters in simulation, we found out empirically out that the fraction of loci with non-extinct alleles, 1 – *C_loss_*(∞), depends mostly on a single composite parameter (Fig. 2A-C)

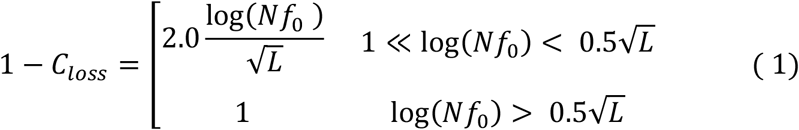

**Fig. 2.**
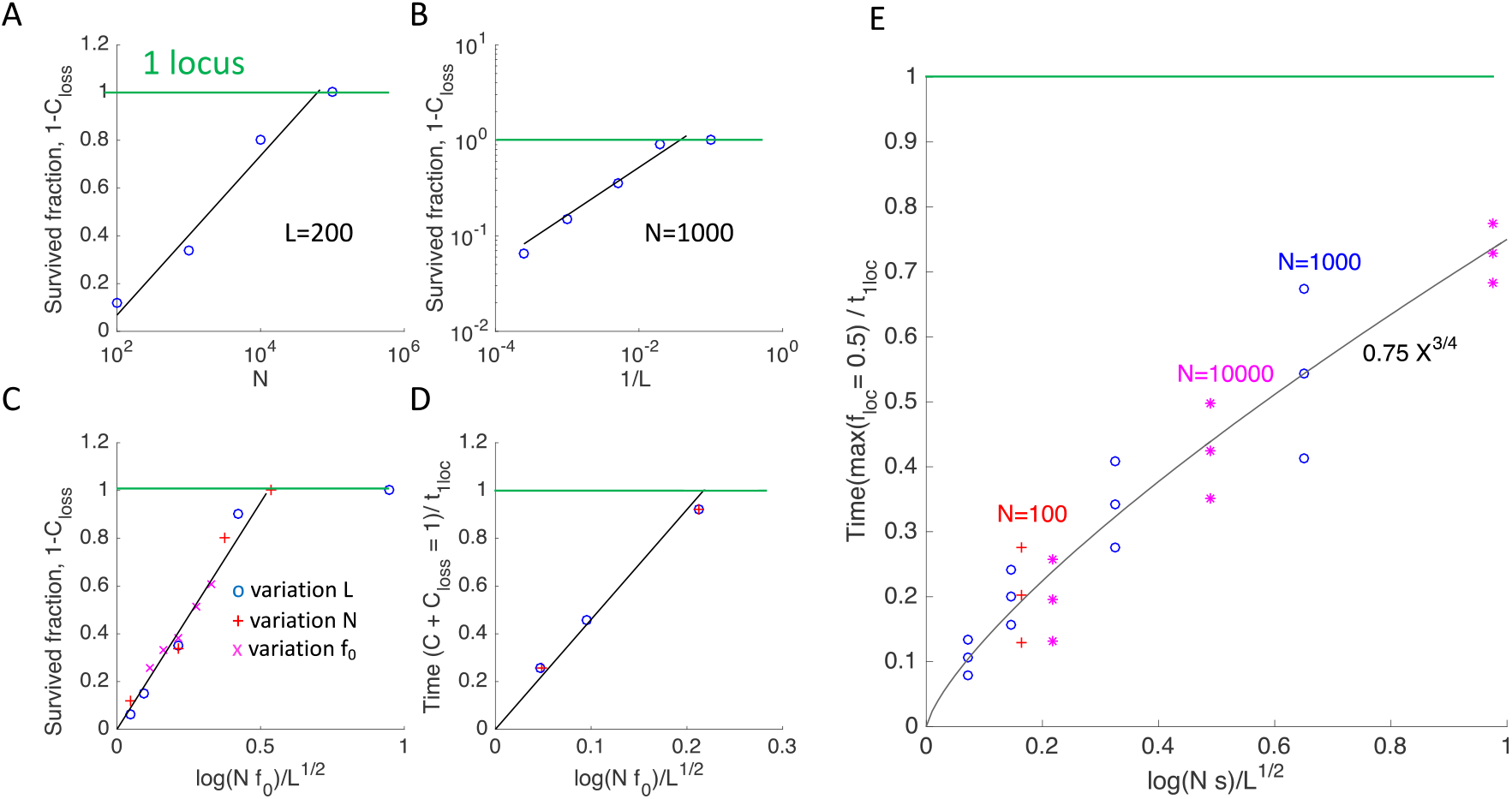
The observables depend mostly on a single composite parameter. **A-C.** The locus fraction where beneficial alleles have survived and completed adaptation, 1 – *C_loss_*(∞), is linearly proportional to the natural logarithm of the population size, log *N*, the inverse square root of the locus number, 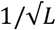, and a composite parameter, 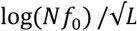. Colored symbols 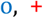, and 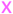 correspond to the variation of model parameters *L, N*, and *f*_0_, respectively, where *f*_0_ > 1/ *Ns*. The green horizontal line shows the prediction of the one-locus model, *C_loss_* ≈ 0. **D.** The time, *t*, when the survived-loci fraction, 1 – *C_loss_*(*t*), equals the average identity by descent, *C*(*t*), [intersection of red and pink curves in Fig. 2] scales linearly with 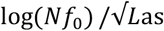 well. **E.** The time when the allelic frequency at the fastest locus reaches 50%, scales as a power ¾ of a similar parameter, 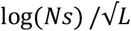. The symbol triplets show the mean and the 95% confidence interval. Colored symbols 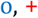, and 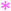 show different values of *N*. The sensitivity to the variation of selection coefficient *s*, crossover number *M*, and initial allele frequency *f*_0_ is shown in S1 Fig and S2 Fig. The default parameter values are *N* = 1000, *L* = 200, *f*_0_ = 0.02 unless shown otherwise. The other parameters are as in Fig. 2.

Note the critical point, 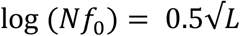. If the population size is too large or the number of loci is too small, no significant loss of polymorphism is predicted.

### The fastest adaptation rate among loci is much faster than in a single-locus model

Because most loci fail to complete adaptation, the average frequency of beneficial alleles per locus, *f_av_*(*t*) saturates far below 1 (Fig. 1A, blue line). The dependence of average heterozygocity on time, 2w^2^(*t*), is decreased accordingly (Fig. 1A, green). The allele frequency averaged over remaining polymorphic sites, *f_pol_*(*t*), increases in the same general time range as the one-locus prediction. The time of half-fixation of polymorphous sites, *t*_50_, is very close to the deterministic one-locus prediction, *t*_50_ ≈ *t*_1*loc*_ (Fig. 1B)

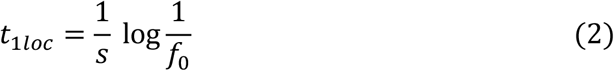

In the range of parameters *s* = 0.025 – 0.2, *L* = 200 – 2000, *N* = 1000 – 10,000, the relative difference between *t*_50_ and *t*_1*loc*_ is between −0.11 and 0.14. Compared to the 1-locus model prediction (blue dashed line in Fig. 1), the dependence *f*(*t*) experiences a delay in the late phases of adaptation and has a noticeable random oscillation component (Fig. 1, blue 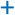).

The speed of adaptation is extremely broadly distributed among loci with non-extinct alleles (Fig. 3H). At some loci, alleles accumulate much faster than predicted by the one-locus model (Fig 1, black line). The half-time of adaptation of the fastest locus, max (*t_loc_*), is much shorter than *t*_1*loc*_ and increases as power ¾ of composite parameter 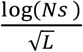 (Fig. 2E) (compare with Eq. 1). The broad variation between loci is created by random recombination events, which bring together different numbers of favorable alleles, and natural selection, which favors the best. As a result, the distribution of genomes in fitness forms a traveling wave well-known for both asexual and sexual populations (40) (Fig. 3A).

**Fig. 3.**
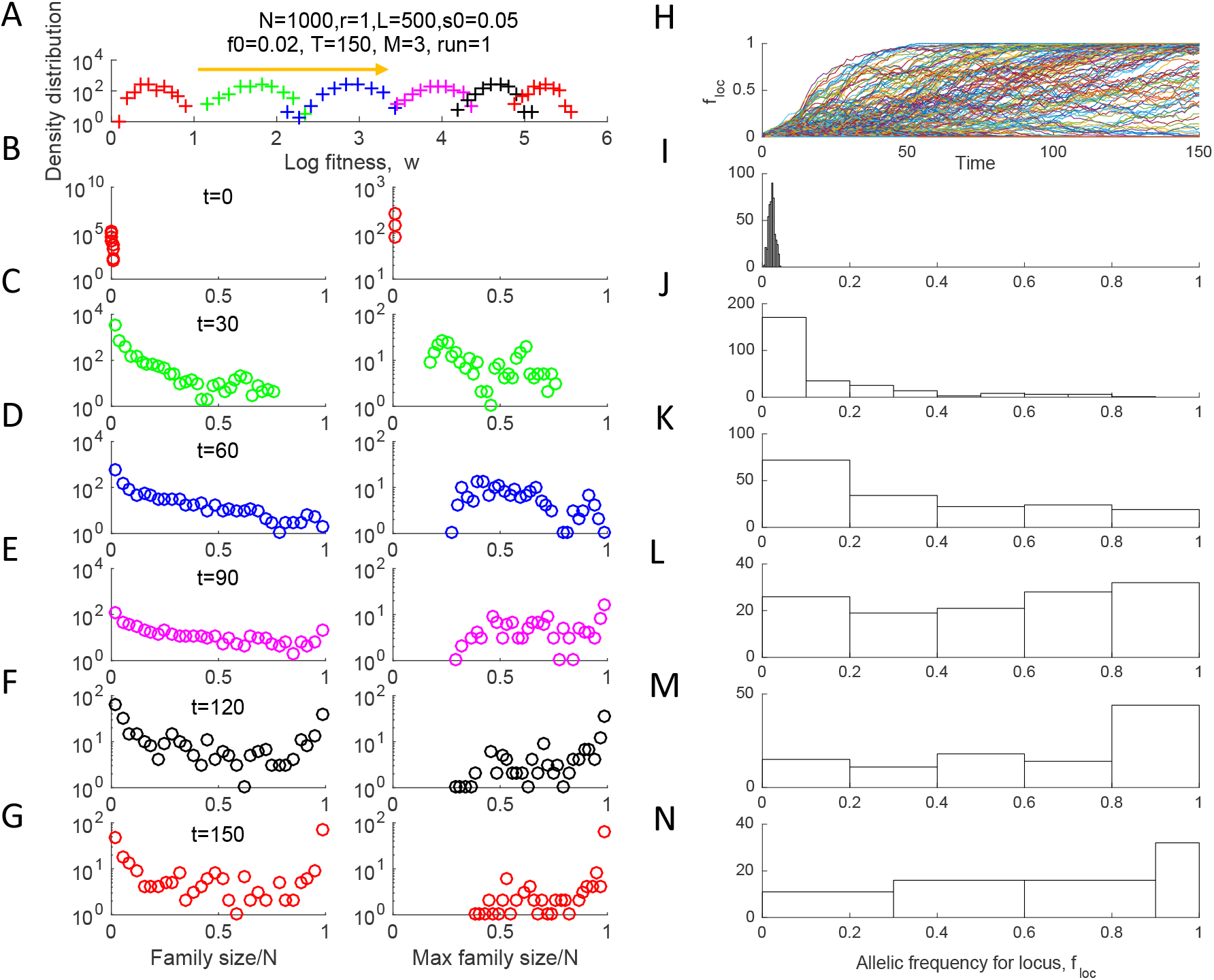
Traveling fitness wave and nonuniform dynamics of separate loci. **A**. Distribution density of genomes in fitness at different time points shown in (B-G). **B-G.** First column: Histograms of the family size defined as the number of sequences with the same initial ancestor at a locus. Second column: Only the largest family per locus is taken into account. **H.** The average allelic frequency for each separate locus, *f_loc_*, as a function of time. **I-N**. Histograms of *f_loc_* across loci at different time points (shown). Parameters are as in Fig. 2.

The fitness classes of the traveling wave have a complex lineage structure that varies between loci. For a given locus, a lineage is determined as the set of individuals that have the same initial ancestor. The lineages all initially consists from a single individual (Fig. 3B), but their sizes grow in time, at different rates for different loci, and become distributed in a very broad range (Fig. 3C-G). The size distribution shifts in time towards larger lineages eventually occupying almost the entire population. If we take into account only the largest lineage for each locus, their size distribution looks similar but has a low cutoff increasing in time (Fig. 3B-G, column 2). The largest lineages grow to a half of the population at a much earlier time than *t*_1*loc*_ in Eq. 2.

### Phylogenetic time scale depends only on the same composite parameter

Another quantity affected by linkage effects is the identity by descent, *C*, defined as the probability of a homologous locus pair to have the same initial ancestor. The average identity by descent averaged over all loci and over only polymorphous loci is almost the same (magenta line and magenta 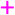, Fig. 1A). This result differs from the single-locus model, where common ancestry is rare, 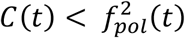, because each of the pair of loci must fall into the same growing lineage to have the same ancestor, and the size of each lineage relative to the population size is smaller than *f_pol_*(*t*). In contrast, in our case, *C* is larger than *f_pol_*(*t*), which is larger than 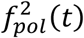. At the time point *T*_2_ where *C* = 1 – *C_loss_*, both values are both close to a half in a broad parameter range, *C*(*T*_2_) ≈ *C*_loss_(*T*_2_) ≈ 0.5. The dependence of *T*_2_ on model parameters can be fit by the formula

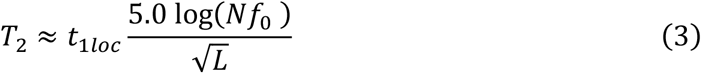

In other words, *T*_2_ is proportional to the same composite parameter that controls the fraction of fixed loci, 1 – *C_loss_*, Eq. 1 (Fig. 3D). Time *T*_2_ defined by Eq. 3 represents a proxy time scale of the phylogenetic tree. Although, at this time point, a population does not have a single ancestor for an average locus as yet, *T*_2_ approximates the time to the most recent common ancestor by an order of magnitude.

### Weak dependence of all observables on the average number of recombination crossovers

The above results in Figs 1 to 3 are weakly sensitive to the average crossover number, *M*. In its entire range of between *1* and *L*, the fraction of loci that do not lose alleles, 1 – *C_loss_*(∞), varies only by the factor of ~2 (Fig. 4).

**Fig. 4.**
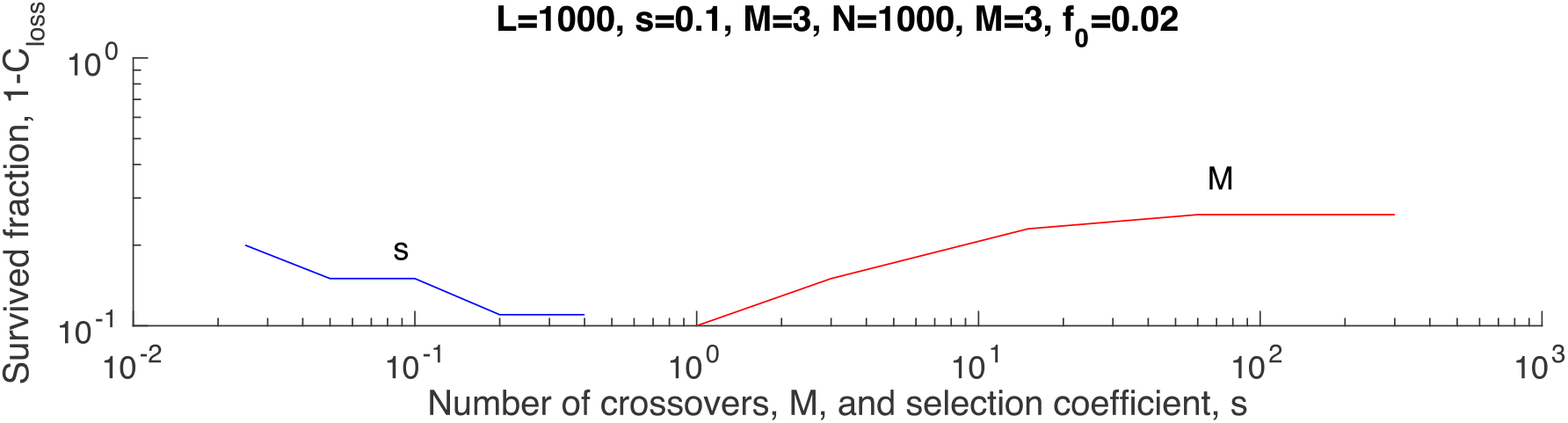
Weak sensitivity of the fraction of loci that complete adaptation to the selection coefficient and the average crossover number. The default parameter values are shown on the top.

### The absence of long-range linkage disequilibrium

No linkage disequilibrium is predicted in the long range. Pearson’s correlator between allelic frequencies at two loci defined as

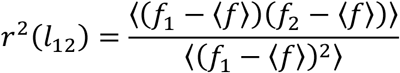

decreases rapidly with the distance between loci, *l*_12_, and the characteristic distance of the decrease shrinks with time (Fig S1). In other words, alleles at far loci segregate independently, as they should in the presence of recombination.

### Far blocks of genome do not evolve independently

The above results for the phylogeny time scale differ from that of scaling theory (57). In my notation, their general result for the average time to the most recent common ancestor has the form [(57), Eq. 5]

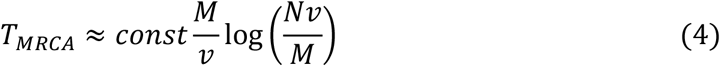

where *v* is the average rate of long-term adaptation, defined as the fitness gain per unit time, *const* is a number on the order of 1, and the logarithm is supposed to be much larger than 1. In my case, the proxy of *T_MRCA_* by the order of magnitude is *T*_2_ in Eq. 3, and the adaptation rate is (see Fig. 1A)

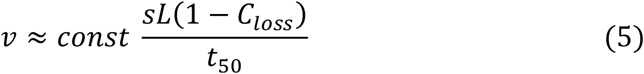

As already mentioned, the average time to a half-fixation for the loci that do not lose alleles, *t*_50_, is always close to one-locus limit *t*_1*loc*_. Substituting Eq. 3 and Eq. 5 into Eq. 4, we get

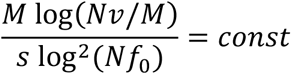

which is clearly false, because M, N, and *s* are independent parameters. Hence, Eq. 4 does not work in the case with pre-existing variation.

Note that the analytic argument in (57) was developed and tested for a different scenario, when the sexual evolution is limited by new mutation events. It was based on two statements: the assumption that a genome evolves as quasi-independent asexual blocks, and an expression for the time to the most recent common ancestor in terms of the average adaptation rate. The expression was based on the basic concept that the time to most recent common ancestor is the lead of the wave divided by the adaptation rate and was confirmed for various multi-locus models, both sexual and asexual. Therefore, it is likely that the quasi-independence assumption is the cause of the discrepancy. In other words, in the case of pre-existing variation, the genome does not evolve as a set of quasi-independent segments. That conclusion is indirectly confirmed by the results in Fig. 3 showing that beneficial alleles can form highly-fit genomes whose rapid growth outruns mixing of genomes due to recombination (Fig. 3). A recombinant that decreases fitness is not relevant for future generations.

Furthermore, within one realization (Monte Carlo run), the fitness variance of a genomic segment normalized to the genome fitness is not linearly proportional to its length, but shows a complex step-like dependence (Fig. 5).

**Fig. 5.**
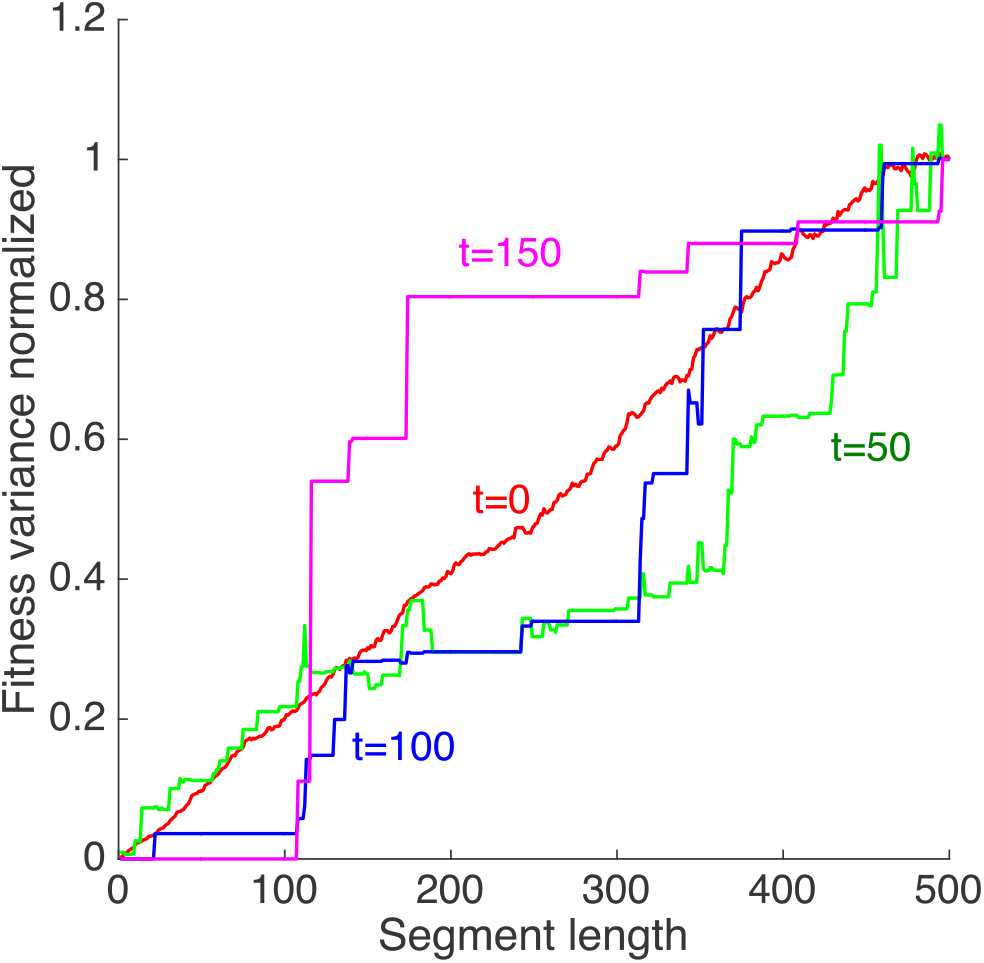
Non-linear dependence of genome segment variance on segment length. X-axis: the length of a genome segment starting from locus 1. Y-axis: Fitness variation between homologous genomic segments divided by the genome fitness variation at the same moment of time. A single run is shown. Parameters are, as in Fig. 1.

### Alleles are fixed inter-dependently

The fixation probability of an allele can be calculated as

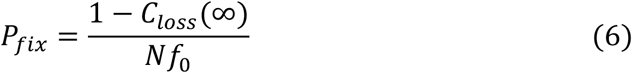

In the parameter interval of interest, this value falls far below the 1-locus prediction, 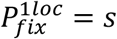 (Fig S2). Probability *P_fix_* plateaus on the value of *s* in the dilute limit of sufficiently small *f*_0_, which agrees with a previous finding in the case of rare mutations, see the limit *r* ≫ *s* in (30). Based on simulation, the transition point to the dilute limit 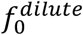 decreases with *N* and *Ĺ*. One can determine the transition point from condition 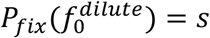 and Eqs. 1 and 6. Replacing log(*Nf*_0_) with 1 if it smaller than 1, we obtain

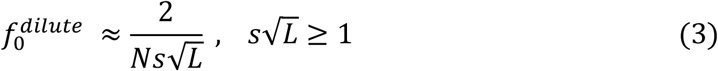

This estimate agrees with the simulation results in Fig S2.

### Phylogenetic tree and allele surfing

In addition to calculating the phylogeny time scale (Fig. 2D), we constructed the ancestral trajectory of a locus between individuals in real-time by memorizing the parentage of each individual locus and then tracing its ancestry back in time. Lineage of each locus jumps among individuals randomly due to recombination (Fig. 6A). If we straighten these trajectories and keep only the topology of coalescence and the coalescent times, we arrive at phylogenetic trees for different loci (Fig. 6B-D). As expected, the tree varies strongly across loci, and the early branches are relatively shorter than in the neutral Kingman’s coalescent. The average density of coalescent events averaged over 10 runs and normalized to the prediction of the selectively-neutral model (*Methods*) decreases exponentially with time (Fig. 6E, F), as it would also in the one-locus limit. This is because coalescent density is proportional to the inverse effective population size (58), which is the size of the growing variant subpopulation. However, the coalescent density is also much larger than in the one-locus limit and increases with number of loci L. Thus, in agreement with the previous studies for various models, uncompensated linkage in the presence of selection makes phylogenetic trees denser and changes their shape by making early branches shorter (24, 27, 56, 59) (Fig. 6E, F).

**Fig. 6.**
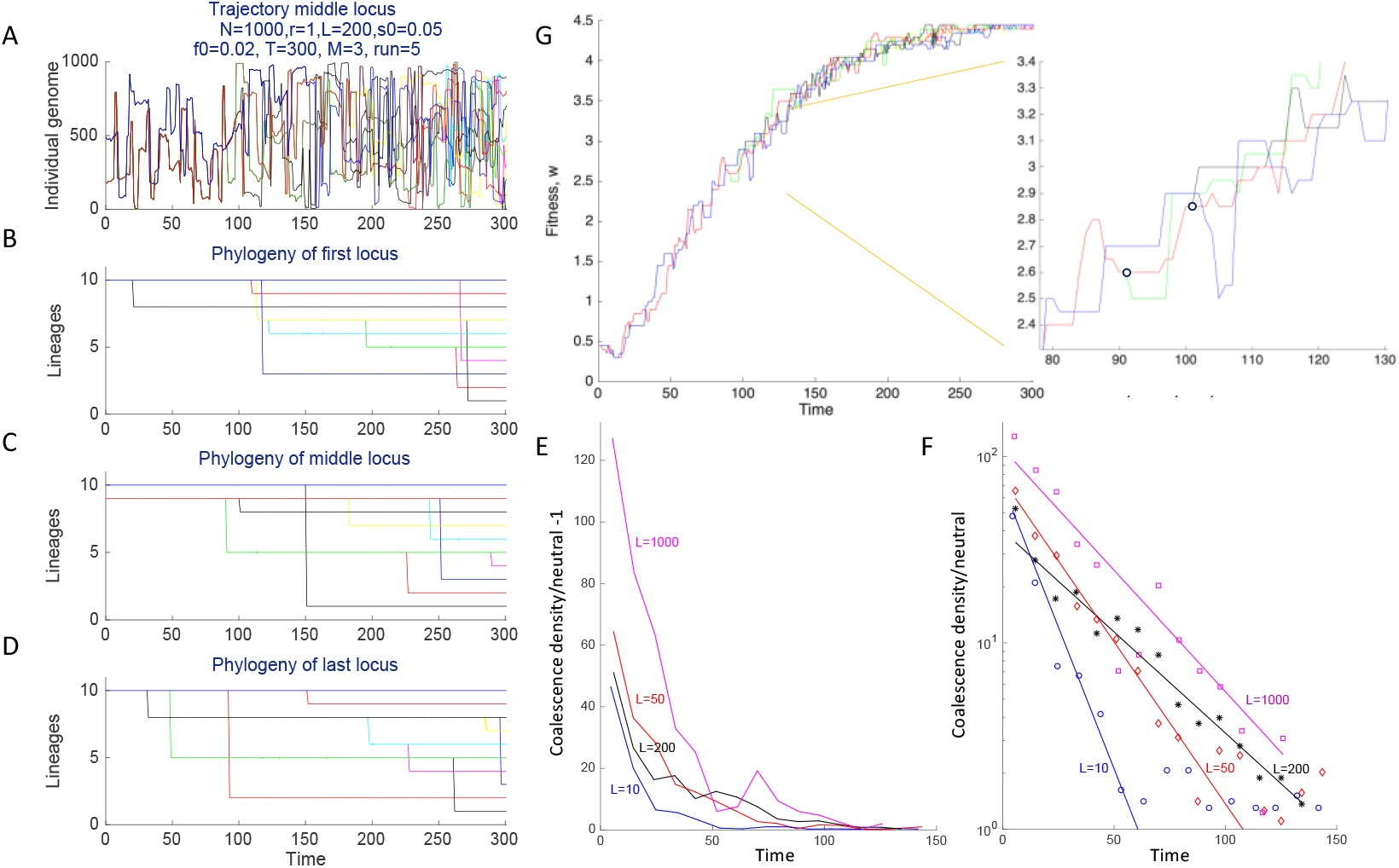
Phylogenetic tree and ancestral history of separate loci. **A.** A reverse-time trajectory of the middle locus (*i* = *L*/2) in 10 individuals numbered 1, 101, 201,… 901 at time *t* = 300 obtained by tracing ancestral history. **B-D.** Phylogenetic trees for three loci (first, middle, and last). **E-F.** The time density of coalescent events averaged over 10 simulation runs and normalized to their values predicted by the selectively neutral model (*Methods*). Linear (E) and logarithmic (F) scales are used for Y–axis. **G.** Fitness trajectories for the middle locus in (A, C). Insert: a small segment is shown by the orange square. Parameters are on (A) unless shown otherwise.

In addition to the trajectory of a locus over specific ancestors (Fig 6A), we can also construct its fitness trajectory, by memorizing the fitness values of its ancestors (Fig 6G). The fitness trajectory comprises alternating straight horizontal segments due to the clonal expansion connected to jumps caused by recombination. The jumps occur in both directions, but more often towards a genetic background with a higher fitness (Fig. 6G).

This “allelic surfing” behavior with vertical and horizontal segments was predicted analytically for sexual populations with a small outcrossing rate (30, 60).

## Discussion

I modeled numerically stochastic sexual evolution of a multi-locus system due to natural selection and pre-existing variation in the form of small numbers of beneficial alleles. Despite of the lack of observable LD for far-situated loci, simulation predicts the existence of string long-range linkage effects encompassing the entire genome. The effects include the extinction of beneficial alleles at most loci due to clonal interference, weak sensitivity of most observables to the average number of crossovers, and a very fast evolution at a fraction of loci. These results are in striking contrast to the previous findings for the long-term evolution driven by mutation, selection, and recombination, where genome was demonstrated to consist from quasi-independent blocks (57). The linkage effects are predicted only for sufficiently long genomes.

If the locus number is decreased, or if the population size is increased, a transition to the independent-locus limit is predicted. The predicted dependence of all linkage effects on the population size is logarithmic (Fig. 2). For a genome of 200 loci and *f*_0_ = 0.02, *s* = 0.1, the transition to the independent-locus regime can be observed already for 100,000 individuals. For a longer genome of 1000 loci, however, loci evolve independently only for populations of 10^12^ individuals or larger, which is unrealistic for most species. A human or an animal population has millions of variable loci, of which a sizeable portion is under selection, so that independent-locus models, probably, never work in most animals, except for rare mutations that are under very strong selection pressure.

We have investigated the case of a constant selection coefficient, but the results are expected to apply also for a sufficiently fast decaying distribution of selection coefficients, such as a Gaussian distribution. Distributions with long tails may have different properties, where the traveling wave is replaced by pairwise clonal interference (26). The case of an exponential distribution can have a mixed behavior, depending on parameter values (26). The exponential distribution is often observed in experiments on pathogens which fact has been explained in a recent work (34).

The results obtained are directly relevant for the viruses that have frequent recombination, such as HIV, polio, or SARS CoV-2. Similar to seasonal human coronaviruses or influenza virus, SARS-CoV-2 is constantly acquiring new mutations in its genome. Evolution is especially fast in receptor Spike protein (61–64). Two major reasons account for the high speed of evolution, as follows. Firstly, Spike has receptor-binding motives that affect transmission, and their evolution leads to the emergence of VOCs with enhanced transmissibility. Secondly, Spike contains epitopes, regions that are very important for the immune response because of their involvement in binding of antibodies that can neutralize virus. Mutations in epitopes are a major factor that limits the virus recognition by the immune system and, hence, the durability of protection (65, 66).

An important puzzle important for devising future vaccination strategies is the origin of the VOCs produced by large groups of new mutations that emerge all together at once (67, 68). Alternative theories of the emergence of VOCs (69) include reverse zoonosis, the evolution within immunocompromised patients (70–72), and the evolution in population pockets not covered by the genetic surveillance. Still another possibility is the fitness valley effect, a cascade emergence of compensating mutations following a primary mutation inferred for HIV and influenza (33, 73).

Based on the present study, we may add yet another possible explanation. While in another respiratory virus, influenza, we observe only rare reassortment of its eight chromosomes, SARS CoV-2, with its single-chromosome genome, has observable crossover recombination (74–78). Hence, the large packages of mutations may emerge due to the combined effects of recombination and natural selection and represent the sequences formed by the fastest loci (Fig. 3). To understand the importance of recombination for SARS CoV-2, we need to know the frequency of co-infected individuals among all the infected, which determines outcrossing probability r, an important input parameter entering the models of sexual populations (13, 19, 30, 57, 60, 79). For fully sexual reproduction considered in the present work, by the definition, *r = 1*. The outcrossing number for SARS-CoV-2 is presently unknown. It could be quite large due to the possibility of a co-infection during superspreading events (80–83). Methods developed previously to quantify recombination from RNA sequence data for HIV could be re-applied to SARS-CoV-2 (13).

### Conclusion

In sexual populations with pre-existing beneficial alleles, in an exponentially broad range of population size, recombination cannot suppress long-range linkage effects, such as the excessive loss of beneficial alleles at most loci, the lack of dependence on the crossover number, and superfast evolution at some loci. These findings may be relevant for interpreting the emergence of new strains of SARS CoV-2.

## Materials and methods

Consider a fully sexual population with *L* loci comprised of *N* individual genomes. Each locus has initially *Nf*_0_ alleles, 1/*Ns* ≪ *f*_0_ ≪ 1, with fitness benefit *s* ≪ 1. In each generation step, each genome undergoes random crossovers with another, randomly chosen genome, with average crossover number *M* producing a recombinant genome. One of the two parents is replaced with the recombinant. Genome number *j* with *k_j_* favorable alleles is replaced with a random number of its copies distributed according to the polynomial distribution implemented by “broken stick” method, as follows. *N* random points are generated uniformly within the interval [0,N] broken into *N* segments. The length of segment *j* is proportional to the fitness of the corresponding genome *w_j_*

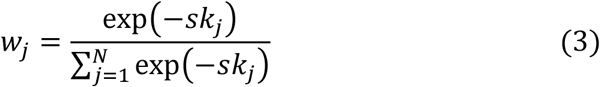

The number of random values that fall into segment *j* are taken to be the number of his progeny in the next generation. Thus, the total number of genomes stays constant. New mutations are neglected, which is shown to be correct in the short-term in the presence of pre-existing genetic variation, both in simulation and experimentally (84). Epistasis is absent. For the modeling studies of epistatic effect, the reader is referred to (31–33, 47–51).

Input model parameters are the selection coefficient across loci, *s* = s_0_, population size *N*, outcrossing rate *r* = 1, number of loci *L*, initial beneficial allele frequency *f*_0_, total simulation time t, average number of recombination crossovers *M*, and the seed number of the generator of pseudorandom numbers.

Parameter ranges studied are *s* = [0.025,0.4], *L* = [10,4000], *N* = [10^2^,10^5^], *M* = [1,300], *f*_0_ = [0.0001,0.02]. The main focus is on the interval of *f*_0_ such that 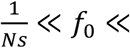. The transition to dilute limit *Nf*_0_*s* ≪ 1 when alleles are fixed independently is shown in Fig. S2.

## Acknowedgement

The study was carried out within the framework of the state assignment of the Federal Agency for Scientific Organizations (FASO Russia: topic no. AAAA-A18-118012290142-9).

## Data and materials availability

The simulation code is available at https://github.com/irouzine/Strong-linkage-in-sex.

## Supporting Information

**S1 Fig.**
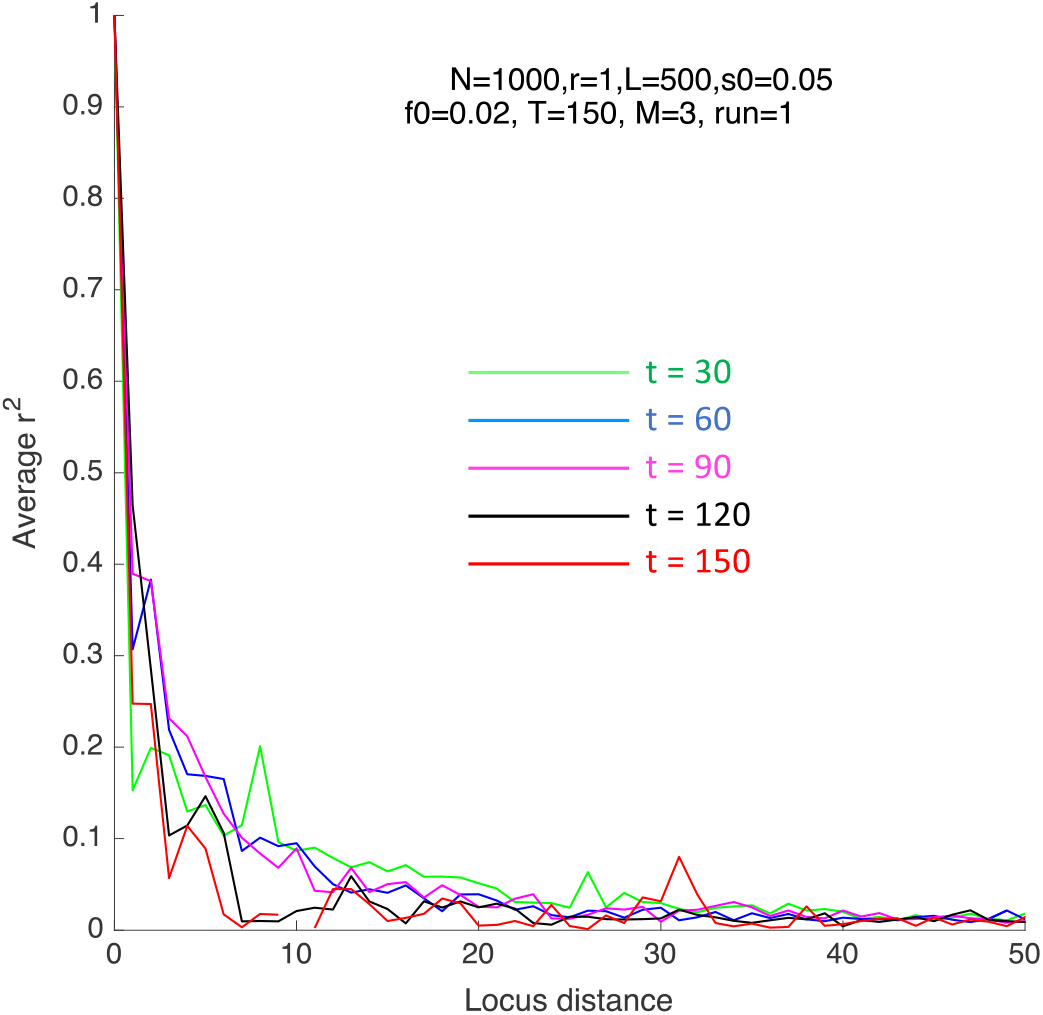
Linkage disequilibrium as a function of distance between loci in the genome. Pearson’s measure r^2^ is averaged over pairs of sufficiently heterozygous loci, 2*f_loc_*(1 – *f_loc_*) > 0.1. The time points and parameters (shown) are the same as in Fig. 3. At *t* = 0, linkage disequilibrium is identically zero due to the initial random distribution of alleles.

**S2 Fig.**
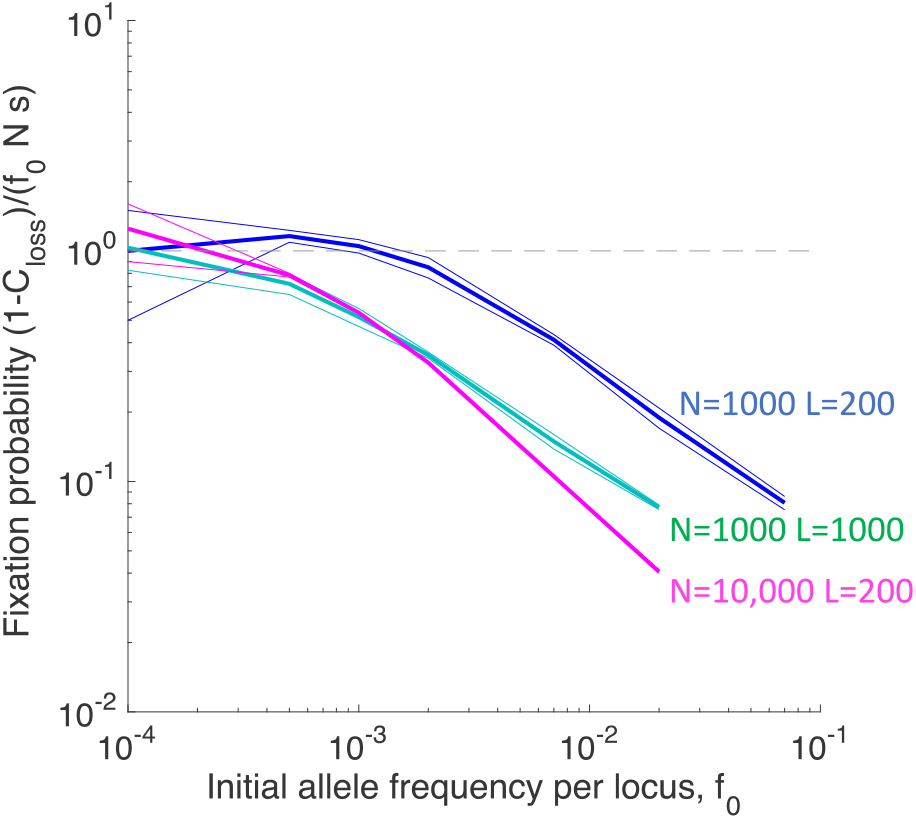
Fixation probability per beneficial allele as a function of the initial allelic frequency exhibits transition to the dilute limit of independent alleles with fixation probability s. Y-axis: The average fraction of surviving polymorphic loci, 1 – *C_loss_*(∞), divided by *f*_0_*Ns*, which is the product of the average number of beneficial alleles per locus, *f*_0_*N*, and the allelic fixation probability in the 1-locus model, *s*. X-axis: The initial frequency of beneficial alleles, *f*_0_. The dependence is shown at three combinations of values of *N* and *L*. Three lines of each color shxow the mean and the mean plus minus the standard deviation between three simulation runs, i.e., the 67% confidence interval. The independent-locus limit of fixation probability shown by the dashed horizontal line is reached at small *f*_0_. Fixed parameters are *M* = 3 and *s* = 0.1.

